# Delineating inter- and intra-antibody repertoire evolution with AntibodyForests

**DOI:** 10.1101/2025.03.11.642619

**Authors:** Daphne van Ginneken, Valentijn Tromp, Lucas Stalder, Tudor-Stefan Cotet, Sophie Bakker, Anamay Samant, Sai T. Reddy, Alexander Yermanos

## Abstract

**Motivation:** The rapid advancements in immune repertoire sequencing, powered by single-cell technologies and artificial intelligence, have created unprecedented opportunities to study B cell evolution at a novel scale and resolution. However, fully leveraging these data requires specialized software capable of performing inter- and intra-repertoire analyses to unravel the complex dynamics of B cell repertoire evolution during immune responses.

**Results:** Here, we present AntibodyForests, software to infer B cell lineages, quantify inter- and intra-antibody repertoire evolution, and analyze somatic hypermutation using protein language models and protein structure.

**Availability and implementation:** This R package is available on CRAN (1) and Github at https://github.com/alexyermanos/AntibodyForests, a vignette is available at https://cran.case.edu/web/packages/AntibodyForests/vignettes/AntibodyForests_vignette.html

## Introduction

Rapid progress in immune repertoire sequencing and artificial intelligence are advancing the field by providing high-quality datasets at single-cell resolution and pre-trained large protein language models (PLMs). These datasets of paired heavy- and light-chain sequences can additionally include informative labels on cellular phenotype, antigen-binding and specificity, as well as protein structure. Moreover, the integration of bulk RNA sequencing data can significantly enhance the resolution of immune repertoire analyses and reduce undersampling issues common to single-cell experiments. Pre-trained PLMs have demonstrated the ability to understand structural and functional properties from protein sequences and have been used to predict features of antibody function (2–7). The modalities of this data play a critical role in unraveling the evolutionary processes of B cells during immune responses. Numerous analyses and tools focus on inferring and quantifying individual antibody lineages (8–15), but there is a lack of software dedicated to studying antibody sequence and structural evolution at the repertoire level. In response to this gap, we introduce AntibodyForests, a comprehensive software designed to thoroughly investigate and quantify the inter- and intra-antibody repertoire evolution (**Figure 1**). AntibodyForests integrates pipelines for analyzing sequence data, associated metadata, phylogenetic tree structures, protein conformations, and PLM features. It introduces novel capabilities beyond existing tools for antibody repertoire phylogenetic analysis, while maintaining comparable computational efficiency (8,11,13–15) (**Table S1, S2**).

**Figure 1.**
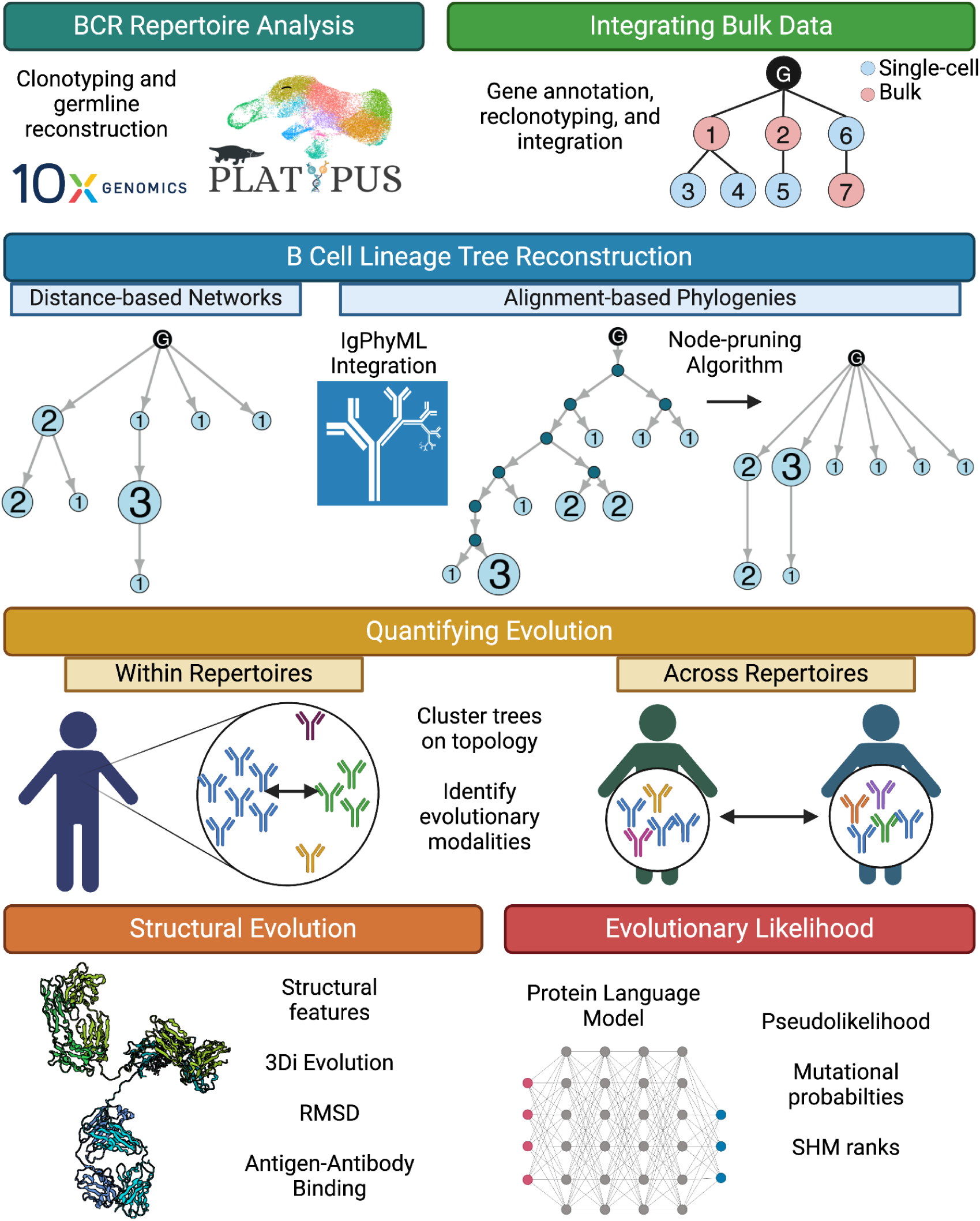
Overview of the AntibodyForests package. This R package is currently composed of a pipeline to reconstruct lineage trees from V(D)J sequencing data preprocessed with the Platypus package and compare trees within and across repertoires. Furthermore, it has modalities to integrate bulk RNA sequencing data, features of protein structure, and evolutionary likelihoods generated with protein language models.

## Usage and application

### Tree reconstruction

AntibodyForests infers B cell mutational networks for reconstructed clonal lineages (clonotypes) **(Figure S1)**. These clonotypes are defined as B cells arising from the same V(D)J recombination event that have undergone somatic hypermutation (SHM) relative to an unmutated reference germline. In AntibodyForests, each clonal lineage is represented as a graph in which the nodes refer to unique antibody sequences, and the edges separating the nodes dictate the clonal relationship between variants. These nodes can be full-length V(D)J sequences, or only certain gene segments or complementarity-determining regions (CDR) from the heavy and/or light chain.

AntibodyForests contains several tree construction algorithms based on either distance matrices or multiple sequence alignments and incorporates minimum spanning tree algorithms and phylogenetic methods. For the distance-based algorithm, AntibodyForests starts by creating a distance matrix based on a user-supplied sequence distance metric. Next, AntibodyForests can create a germline-rooted minimum spanning tree or neighbor-joining tree. Alternatively, the default algorithm of AntibodyForests starts with the germline node and iteratively adds nodes with the smallest distance, comparable with previously developed algorithms (8). In the case of a tie during the default network construction, AntibodyForests has several options that allow the user to select whether breadth or depth (i.e., whether new sequences are added to the node with the most or least descendants), mutational load (whether they are added to the node closest or furthest from the germline), clonal expansion (whether they are added to the node with the highest or lowest cell count), or random addition dictates network topology. For the phylogenetic algorithms, AntibodyForests starts by creating a multiple sequence alignment. Next, AntibodyForests can construct a maximum parsimony tree, minimizing the number of mutations, or a maximum likelihood tree based on various evolutionary substitution models. Alternatively, users can supply previously computed phylogenetic trees in Newick format, thereby enabling the integration of different mutational models, such as those developed specifically for antibodies (9,16,17). To maximize flexibility for the user, AntibodyForests is additionally compatible with the output format of the commonly used IgPhyML tool (18). AntibodyForests can further interrogate and visualize network robustness by comparing tree topology across different tree construction methods using metrics such as the generalized branch length distance (GBLD) (19) **(Figure S4)**. Although the package is designed for single-cell immune repertoire data, the pipeline can also be used with bulk repertoire sequences and even has a function to integrate the two data modalities **(Figure S3).**

Our framework diverges from the traditional B cell phylogenetic tree structure by enabling recovered sequences to serve as either internal or terminal nodes, in addition to allowing for multifurcation events. Some phylogenetic methods generate internal nodes serving as the most recent common ancestors (MRCA). In antibody repertoires, these MRCAs can be recovered in the sequencing data (8,20,21), resulting in nodes with zero branch length to the internal parent node, or they can represent B cells that are not part of the repertoire at the time of sampling (**Figure S2**). To address this, AntibodyForests offers flexibility of handling internal nodes, allowing users to tailor tree representations to either lineage networks of sampled sequences or phylogenetic trees with unsampled intermediate cells. To transform traditional bifurcating phylogenetic trees into multifurcating lineage networks, AntibodyForests includes various algorithms to remove these inferred internal nodes. For example, users can choose to remove only internal nodes with zero branch length to a terminal node, which preserves mutational ordering. Additionally, internal nodes can be replaced by their closest descendant or directly linked to an upstream parental node, favoring either depth or breadth in tree topology. Removing internal nodes potentially impacts the evolutionary trajectory and thereby influences downstream analyses and interpretations; however, how different methods impact tree topology can be quantified using functions within AntibodyForests. When comparing different trees and repertoires to each other, it is important to use the same method for internal node removal.

Upon completion of the network, the color of each node can correspond to the fraction of cells with a given isotype, transcriptional cluster (if such information is available), or a custom BCR feature **(Figure S2)**. To highlight receptor sequences that are identical across multiple B cells, the node size can be scaled to match the relative clonal expansion, while the label can be adjusted to depict the variant frequency. To understand the evolutionary distance separating two nodes, it is possible to include edge labels that correspond to the distance separating nodes.

### Quantifying intra- and inter-antibody repertoire evolution

A quantitative interpretation of antibody lineage trees is crucial to identify different patterns of clonal selection and expansion that drive B cell evolution within and across repertoires. A repertoire-wide analysis of these tree topologies can identify features of SHM and could identify potentially interesting therapeutically relevant antibodies or vaccine targets. For example, Wu et al. discovered a broadly neutralizing antibody against HIV-1 by identifying progenitor sequences preceding viral neutralization capabilities using phylogenetics (22).

Phylogenetic methods have been extensively used to investigate single B cell clonal lineages (12,22–26). However, the evolution of the repertoire as a system has not been fully explored using the collection of lineage trees. AntibodyForests includes two modules explicitly dedicated to comparing evolution within and across repertoires, in addition to comparing topologies consisting of identical sequences **(Figure S4).** Tree topology metrics can be either calculated and stored within the AntibodyForests object or into a separate matrix that can be supplied for downstream clustering and visualization. We have included various metrics to describe tree topologies, including general graph theory metrics and more specialized metrics. For example, we included the Sackin index for tree imbalance, where a high index represents longer branches and more nodes arising from a specific descendant and thereby suggests the presence of selective pressure (27). Users can also compute various properties of the Laplacian spectral density of the lineage trees. These properties characterize specific evolutionary patterns such as species richness (principal eigenvalue), deep or shallow branching events (asymmetry), and tree imbalance (peakedness) (28). Together, AntibodyForests’ metrics can be used to project topology features using dimensionality reduction techniques and clustering, thereby identifying topologically similar networks. Clusters of trees can be related to features specific to single-cell immune repertoire sequencing, such as transcriptional phenotype, isotype, or expansion, as well as other custom metadata features provided by the user.

To illustrate the suitability of AntibodyForests to comprehensively reconstruct and analyze whole BCR repertoires and study patterns of SHM upon immune activation, we included example analyses in the supplementary material using single-cell BCR data collected from the blood and lymph nodes of individuals post SARS-CoV-2 vaccination (24). Using this dataset, we demonstrate example analyses and output plots that arise directly from functions internal to AntibodyForests while also confirming results present in the original paper **(Figures S5A-E)**. This includes comparing tree topologies both within and across repertoires using the aforementioned metrics, and we further provide examples on how users can seamlessly relate repertoire features such as V-gene usage and isotype to evolutionary features.

### Relating protein language model likelihoods to B cell evolution

PLMs have demonstrated success in understanding features of sequence, structure, and function of natural proteins (general PLMs) (29–32). Following this success, additional PLMs trained specifically on antibody sequences were able to learn B cell specific features such as germline usage and SHM (antibody-specific PLMs) (2–6,33). The learning objective of PLMs allows for the computation of a likelihood score for each residue in the input sequence. These scores can be used to calculate a so-called pseudolikelihood for the entire sequence. PLM-based pseudolikelihoods have been leveraged to capture global evolutionary trajectories (34). Furthermore, general PLMs have been demonstrated to improve antibody affinity through in silico mutagenesis by mutating for increasing evolutionary likelihood (7). Given the relevance of PLMs in studying and engineering antibodies, we included PLM-guided metrics within AntibodyForests to describe B cell evolution at the level of an entire repertoire and a single lineage. Both per-residue likelihoods and per-sequence pseudolikelihoods can be incorporated into AntibodyForests to evaluate correlations between tree topology and PLM-based likelihoods of mutating and conserved residues along the edges of the trees and compare within and across repertoires **(Figure S6A)**. We have provided examples of how AntibodyForests can be used to compare PLM-based likelihoods both within and across repertoires and have further highlighted the types of plots that can be directly produced with functions internal to AntibodyForests (**Figure S6B-E**). Users are able to compare how PLM-based likelihoods change as a function of B cell evolution, which can provide insight into which residues may be more likely to evolve and which amino acids are tolerated during SHM. The computational pipeline present in AntibodyForests can help users perform PLM-guided antibody engineering (7) and further be used to study selection signals of antibody repertoires (5). Importantly, we have ensured that AntibodyForests is compatible with many types of PLMs, including both general and antibody-specific PLMs. Users can avoid potential biases that may be present in antibody databases, such as the high proportion of germline-like antibodies, by leveraging PLMs trained on diverse types of protein sequences that also include filters based on homology thresholds (e.g., ESM family of PLMs (29,31)), by fine-tuning custom PLMs, or by using PLMs explicitly trained to avoid germline biases such as Ablang2 (33).

### Structural evolution

Recent computational efforts have massively accelerated the ability to predict protein structures from sequence alone (31,32,35,36). Although SHM acts on the sequence directly, the structure of the antibody determines binding affinity with antigens. We therefore included a structure-specific module in AntibodyForests to integrate antibody structural evolution (**Figure S7A**). This allows users to quantify structural features as a function of sequence distance from the germline. Examples of these features are: 1) structure similarity (root-mean-square deviation (RMSD)), 2) biophysical properties (hydrophobicity, charge, 3Di alphabet (37), free energy, and pKa (38)), and 3) global confidence of the prediction (predicted Local Distance Difference Test (pLDDT)). Furthermore, we have added the option to include antibody-antigen complexes and calculate how properties at the binding interface change with SHM. In addition to RMSD to the germline, AntibodyForests can calculate a pairwise RMSD between the structures connected with edges in the lineage trees and calculate their correlation with the number of mutations. To illustrate a use case of the AntibodyForests structural functions, we highlight how AntibodyForests can integrate structures from Alphafold3 of antibody-antigen complexes to quantify structural evolution within individual lineages **(Figure S7)**. This functionality enables detailed analyses of how SHM, in response to immune stimuli such as vaccination, may drive structural divergence and functional maturation. Although structural modeling methods still struggle with the complex and variable nature of antibody CDR loops and individual mutations on structural predictions (36,39,40), the framework included in AntibodyForests will remain relevant as deep learning algorithms continue to improve.

### Concluding remarks

Taken together, AntibodyForests can both infer and visualize individual clonal lineages from single-cell V(D)J sequencing data and incorporate data such as bulk V(D)J sequences and transcriptional phenotypes. AntibodyForests can incorporate trees constructed with existing tools, and transform its bifurcating phylogenetic output to a more accurate multifurcating representation of the inferred antibody lineages. The package hosts various algorithms to compare evolution within and across repertoires using graph theory, protein language models, and protein structure. With the ongoing increase in immune repertoire sequencing data and the continuous improvement of deep learning strategies, we believe AntibodyForests is an invaluable toolbox to analyze B cell evolution and selection at both the clonal and repertoire levels.

## Supplementary data

**Table S1.** Comparison of AntibodyForests to other tools for the analysis of B cell lineages. AntibodyForests advances other tools with regard to the downstream analysis of the lineage trees.

|  | AntibodyForests | Dowser<br>(1) | IgTree<br>(2) | SONAR<br>(3) | GCtree<br>(4) | BraCer<br>(5) |
| --- | --- | --- | --- | --- | --- | --- |
| **Lineage reconstruction algorithms** |  |  |  |  |  |  |
| *Distance-based networks* | + | - | + | - | - | + |
| *Maximum Parsimony* | + | + | - | - | + | - |
| *Maximum Likelihood* | + | + | - | + | - | + |
| *B-cell specific substitution model* | + | + | - | + | + | - |
| *Recovered sequences as internal nodes* | + | - | + | - | + | + |
| *Reconstruct intermediate sequences* | - | + | - | + | - | - |
| *Integrate single-cell and bulk sequences* | + | - | - | - | - | - |
| *Create lineage tree plots* | + | + | - | + | + | + |
| **Quantifying sequence evolution** |  |  |  |  |  |  |
| *Topology metrics of trees* | + | - | - | - | - | - |
| *Metadata distribution within trees* | + | + | - | + | - | - |
| *Compare trees based on topology* | + | - | - | - | - | - |
| *Compare (groups of) repertoires based on topology* | + | - | - | - | - | - |
| *Cluster trees on topology* | + | - | - | - | - | - |
| *Analyze longitudinal samples* | + | + | - | + | - | - |
| *Analyze mutations along the edges of the trees* | + | - | + | - | - | - |
| **Quantifying functional evolution** |  |  |  |  |  |  |
| *Integrate protein structure* | + | - | - | - | - | - |
| *Estimated antibody-antigen binding* | + | - | - | - | - | - |
| *Protein Language Model likelihoods* | + | - | - | - | - | - |
| **Accessibility** |  |  |  |  |  |  |
| *Open-source implementation* | + | + | - | + | + | + |
| *Vignette available* | + | + | - | + | + | + |

**Table S2.** Benchmarking AntibodyForest and Dowser(1). Benchmarking runtime and RAM usage of preprocessing CellRanger output and constructing lineage trees for the heavy chain nucleotide sequences of sample SRR17729692 from the Kim et al. dataset (6). *Preprocessing for AntibodyForests uses the Platypus (7) software and Dowser uses Change-O and SHazaM. Preprocessing includes grouping the BCR transcripts into clonotypes, assigning a germline, and preparing a dataframe ready for lineage tree reconstruction. Code available at https://github.com/dvginneken/BenchmarkingAntibodyForests

|  | AntibodyForests | Dowser (1) |
| --- | --- | --- |
| **Preprocessing\*** |  |  |
| *CPU time (seconds)* | **667.71** | 707.31 |
| *RAM usage (megabytes)* | **861.2** | 1,181.6 |
| *Wall-clock time (seconds)* | 708.89 | **169.55** |
| **Maximum Likelihood Tree** |  |  |
| *CPU time (seconds)* | 508.95 | **34.49** |
| *RAM usage (megabytes)* | **738.8** | 914.6 |
| *Wall-clock time (seconds)* | 514.86 | **35.29** |
| **Maximum Parsimony Tree** |  |  |
| *CPU time (seconds)* | **33.47** | 49.31 |
| *RAM usage (megabytes)* | **646.3** | 925.3 |
| *Wall-clock time (seconds)* | **34.04** | 50.93 |
| **IgPhyML Tree** |  |  |
| *CPU time (seconds)* | 587.75 | **1.86** |
| *RAM usage (megabytes)* | 1,076.4 | **24.37** |
| *Wall-clock time (seconds)* | **7,730.09 (2h8m)** | 16,111.44 (4h28m) |
| **Distance-based Network** |  |  |
| *CPU time (seconds)* | **11.54** | *Not available* |
| *RAM usage (megabytes)* | **133.2** | *Not available* |
| *Wall-clock time (seconds)* | **12.14** | *Not available* |

**Figure S1.**
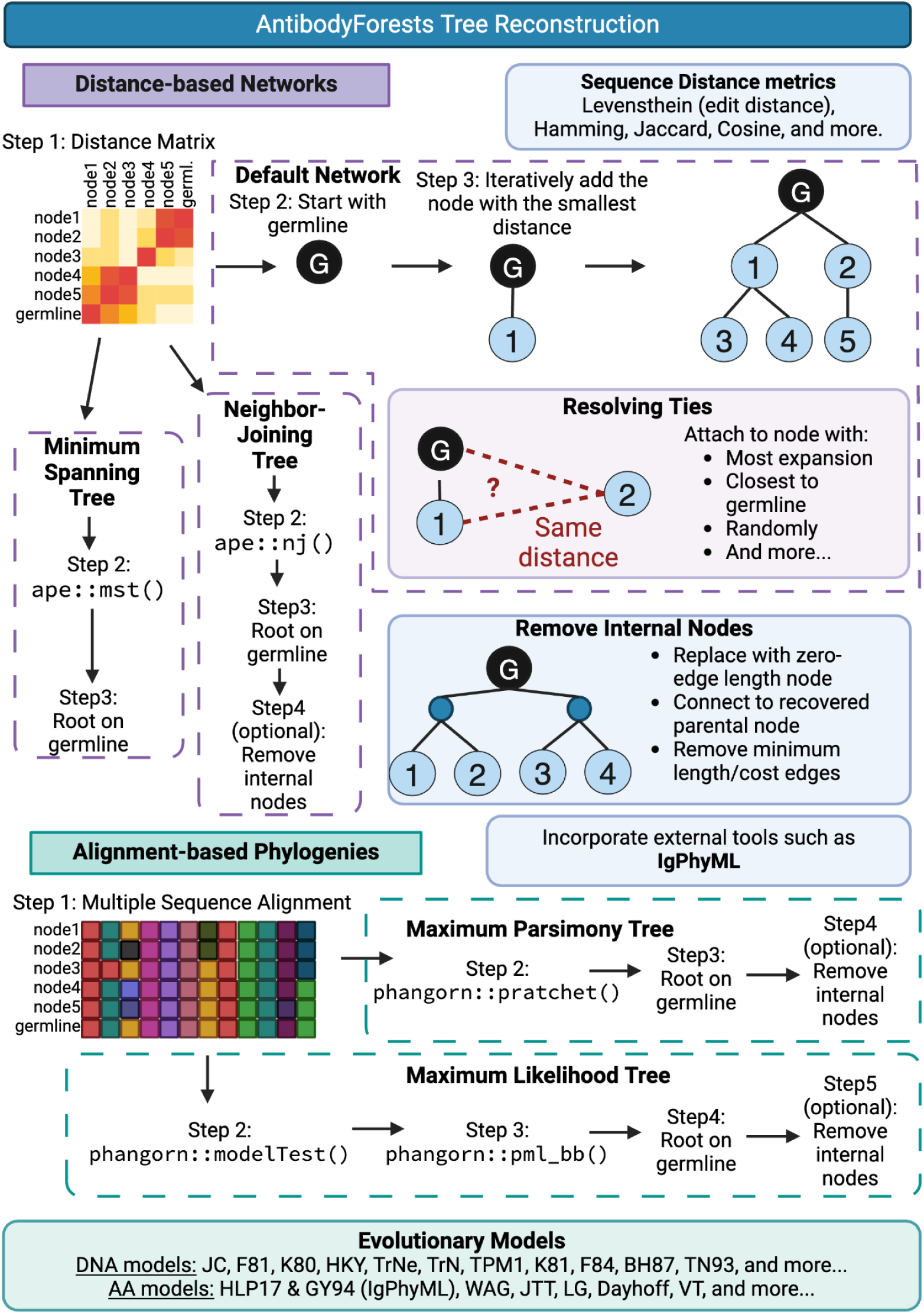
Overview of tree reconstruction algorithms in AntibodyForests. AntibodyForests contains five methods for lineage tree reconstruction and can integrate output from external tools such as IgPhyML (8). AntibodyForests works with various metrics for sequence distance and a wide range of evolutionary substitution models. Additional algorithms are available to resolve ties and remove internal nodes during network construction. Created with biorender.com.

**Figure S2.**
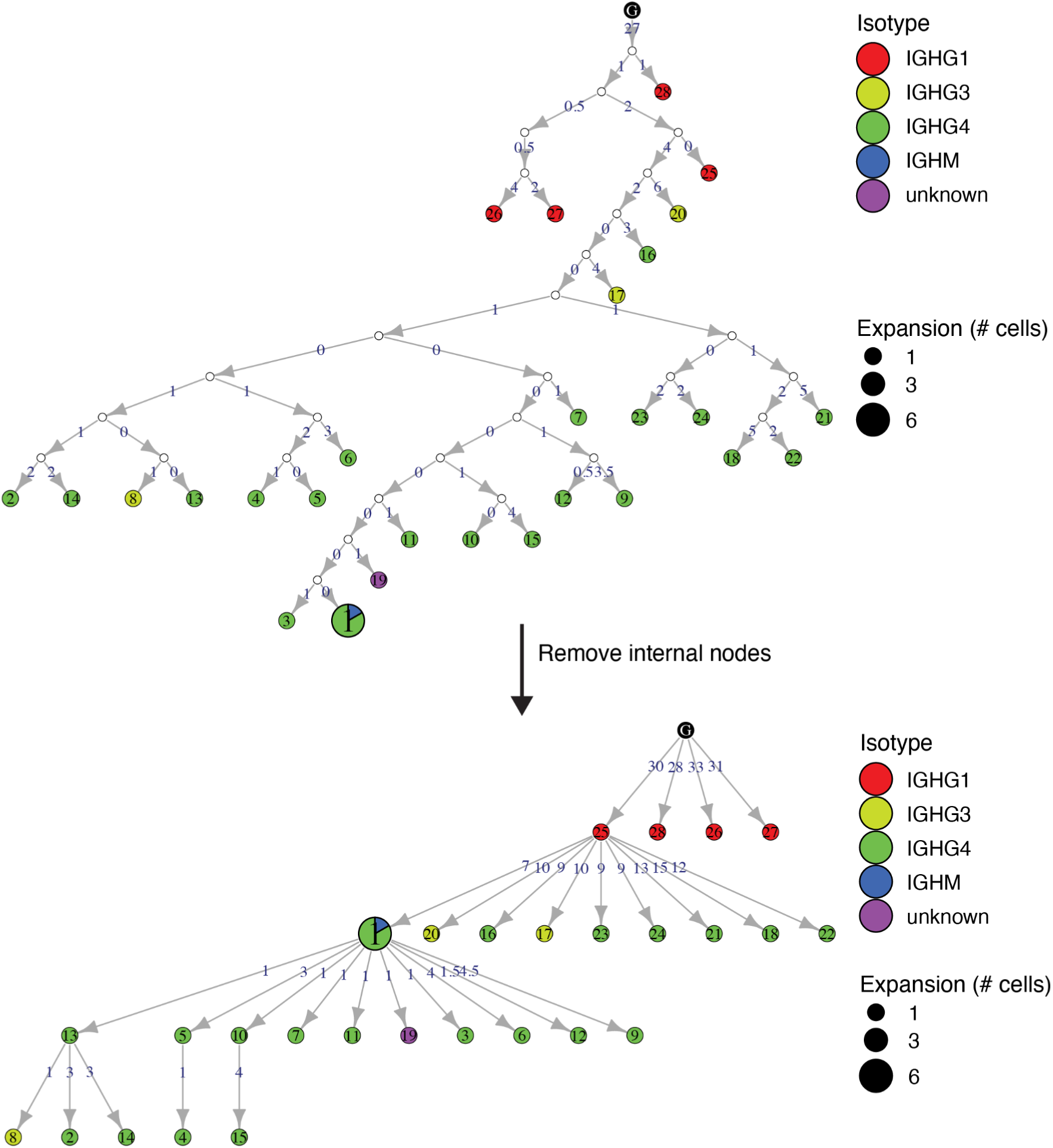
Examples of AntibodyForests trees. An example AntibodyForests object was generated using the maximum parsimony algorithm in the Af_build function. The top tree is before internal node pruning, the bottom tree is after node pruning. The black node on top of the network refers to the germline as indicated by 10x Genomics, the white nodes (top tree) are internal nodes, and the colored nodes are recovered sequences. Node colors correspond to the isotypes. If multiple isotypes are presented within a node, the node is represented as a pie chart. Node size corresponds to the number of cells with that unique heavy+light chain combination. Node labels can be matched to additional information within the AntibodyForests object. Edge labels correspond to the edit distance between the sequence of the attached nodes. Single-cell BCR data for this analysis was derived from Kim et al. (6)

**Figure S3.**
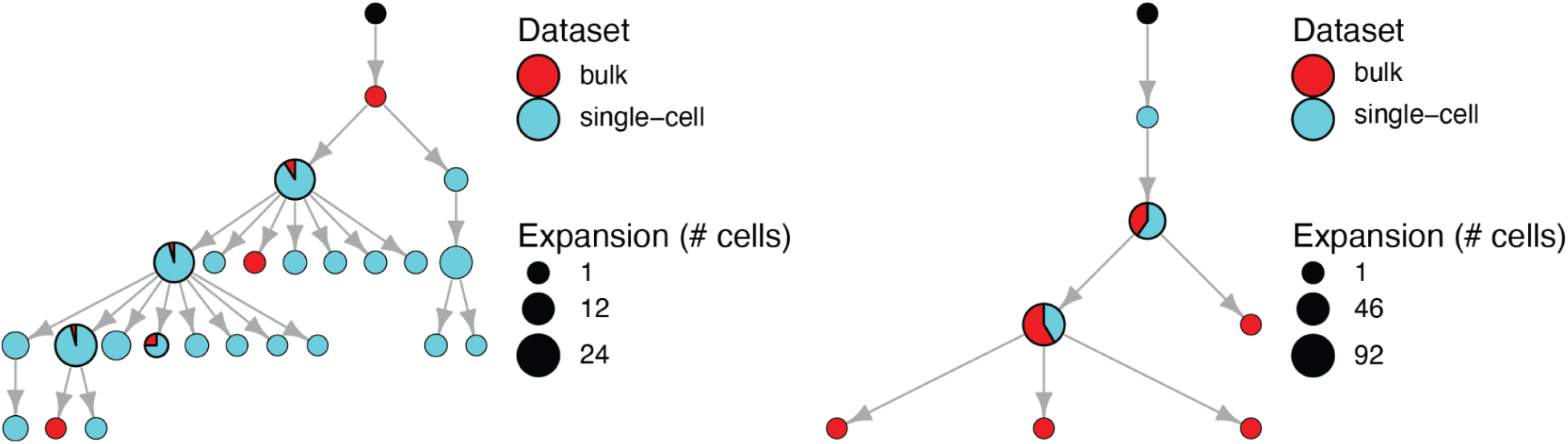
Example bulk RNA integration. An example AntibodyForests object was generated using both single-cell and bulk heavy chain BCR sequencing using the default parameter settings in the VDJ_integrate_bulk and Af_build functions. The black nodes correspond to the reconstructed germlines. These two trees serve as examples to show the identical sequences between the bulk and single-cell datasets (pie chart nodes) and the integration of unique bulk sequences into the single-cell based clonotypes (red nodes). Single-cell and bulk BCR data for this analysis were derived from Neumeier et al. (9).

**Figure S4.**
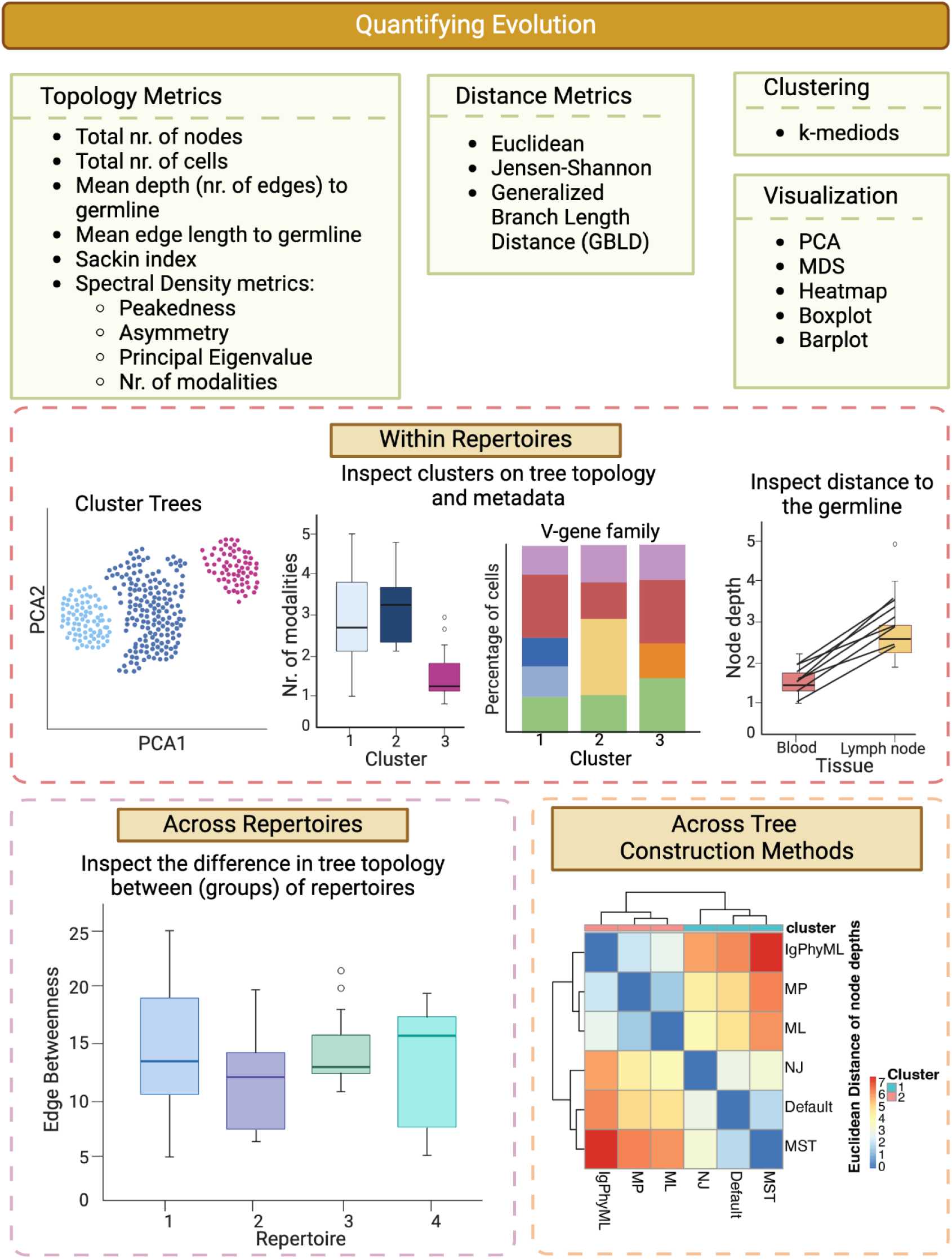
Overview of methods to quantify evolution. AntibodyForests contains metrics to compare within and across repertoires or the same set of trees with different methods. Trees can be clustered based on various distance metrics and visualized in for example dimensionality reduction plots such as a PCA plot. AntibodyForests contains functions to investigate the difference in topology metrics (such as the nr. of spectral density modalities) or metadata of the lineage tree nodes (such as v-gene family) between the clusters. Additionally, distance to the germline (e.g. node depth) can be compared between metadata features of the nodes. AntibodyForests can additionally compare tree topology metrics across repertoires, such as the edge betweenness (the number of shortest paths between two nodes that cross this edge). To compare different tree reconstruction algorithms, AntibodyForests can be used to create heatmaps of distance metrics between the trees, such as the Euclidean distance of node depths. A more detailed explanation of all metrics can be found in the vignette (https://cran.case.edu/web/packages/AntibodyForests/vignettes/AntibodyForests_vignette.html). Created with biorender.com.

**Figure S5.**
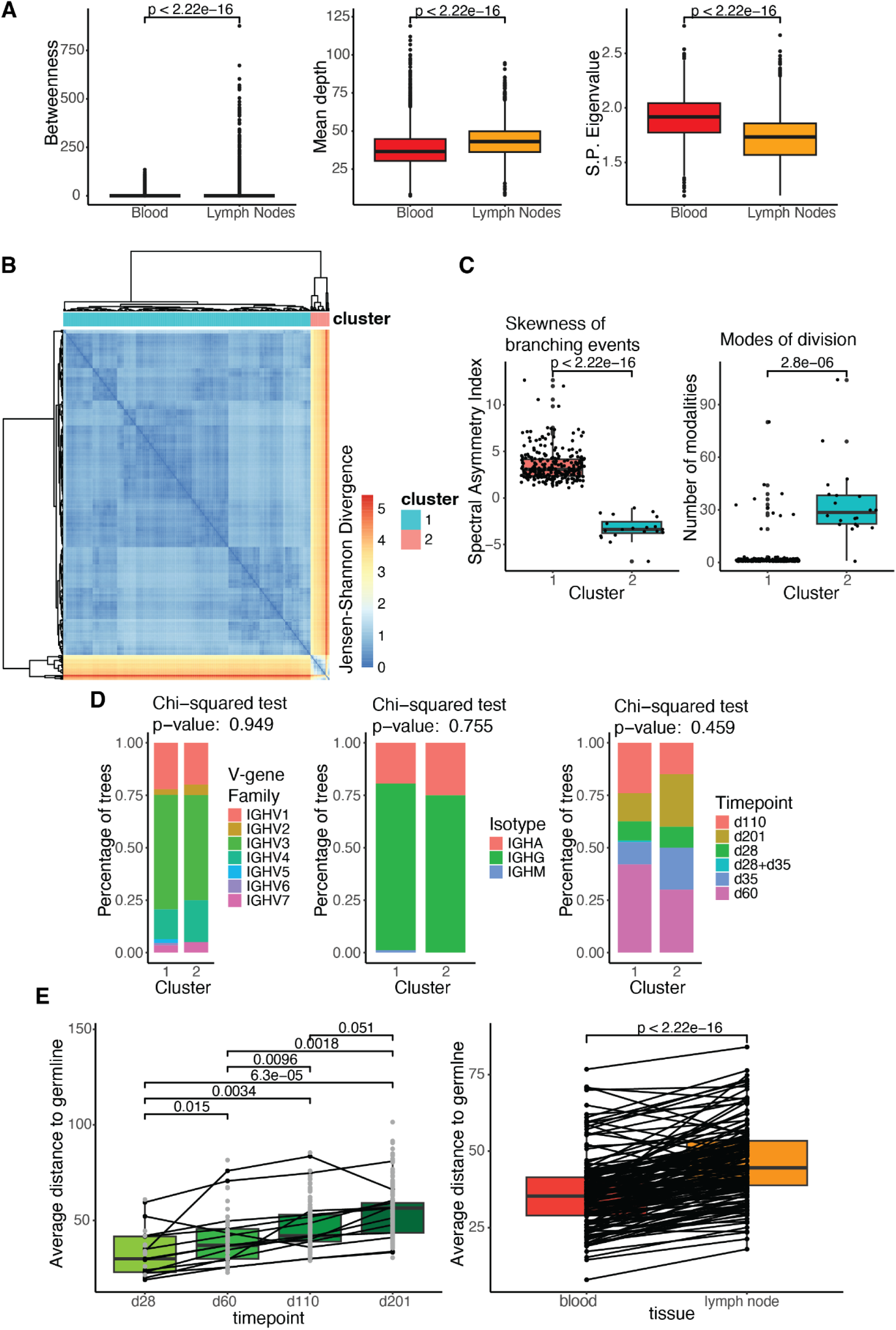
Example of how AntibodyForests can quantify antibody repertoire evolution of single-cell BCR data collected from the blood and lymph nodes of individuals post SARS-CoV-2 vaccination. B cell lineage trees were constructed with the default algorithm using the heavy+light chains using the Af_build function. A) Comparison of tree topology metrics (node betweenness, average edge length to the germline, and spectral principal eigenvalue) between BCR repertoires in the blood and lymph nodes using the Af_compare_across_repertoire function. This across-repertoire comparison revealed the blood repertoire to have more diversity, while the deeper trees in the lymph node repertoire suggest more sequential evolution. B) The B cells from all tissues and time points for each individual were clonotyped together and the trees were clustered based on the Laplacian spectrum. This heatmap shows the Jensen-Shannon divergence in Laplacian spectral density between trees using the Af_compare_within_repertoires function. C) Spectral density metrics of trees in both clusters using the Af_cluster_metrics function. This within-repertoire comparison demonstrated a subset of trees displaying deep branching events, suggesting that cells with a small degree of SHM were recovered. This same subset was characterized by multiple topology modalities, indicating various diversification events. D) Proportion of trees in each cluster and their predominant labels (V-gene family, isotype, and timepoint of sampling) using the Af_cluster_node_features function. E) Average distance (sum of edge lengths) from the nodes of each group to the germline using the Af_distance_boxplot function. Grey dots represent trees not containing all groups. This analysis revealed that SHM increased with the time after vaccination and that blood-derived BCRs were located closer to the germline than those from lymph nodes. Single-cell BCR data for this analysis was derived from Kim et al. (6)

**Figure S6.**
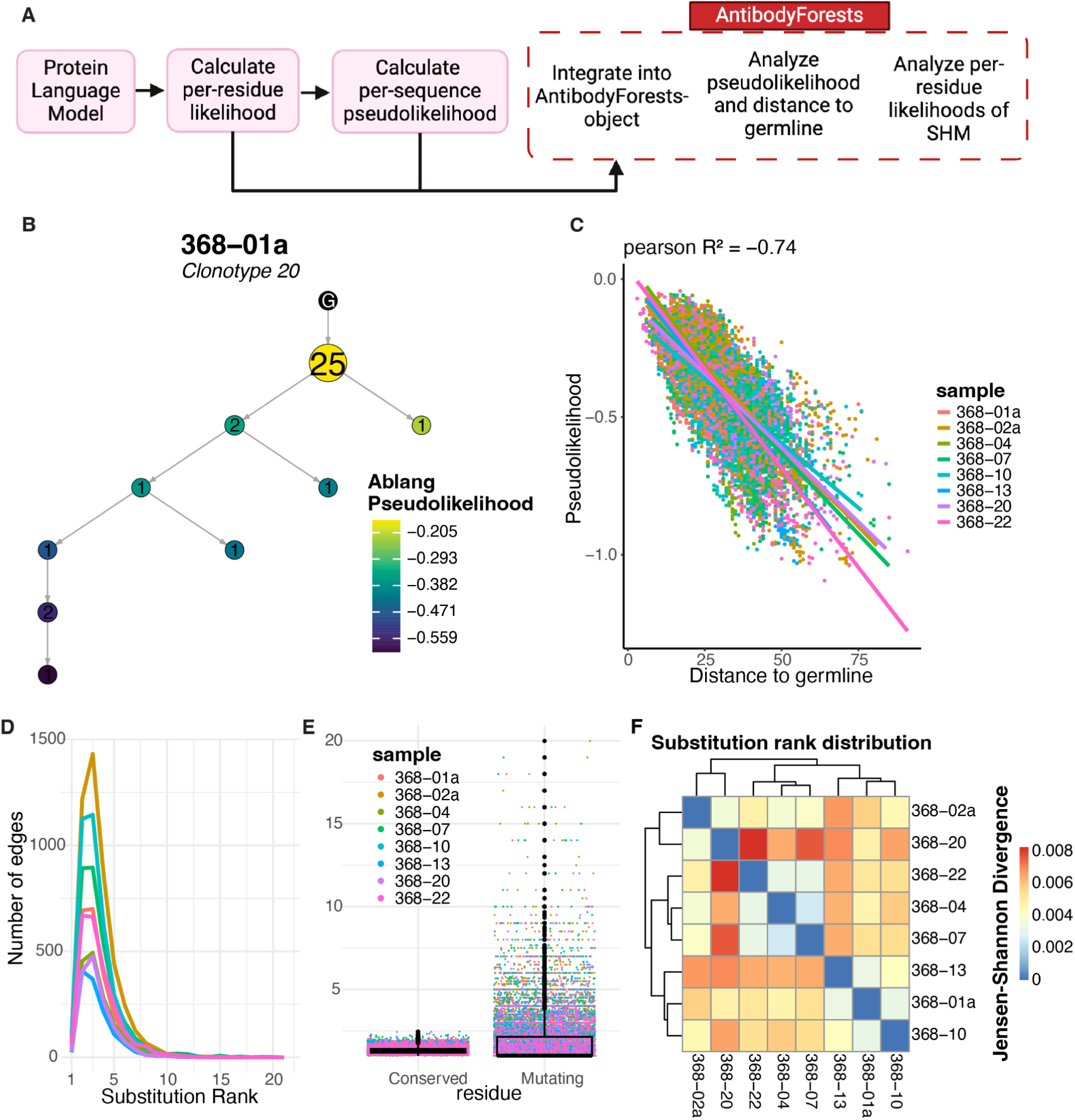
Example of how AntibodyForests can integrate B cell evolution with protein language model (PLM) likelihoods. A) Overview of the PLM workflow integrated with AntibodyForests. After calculating the PLM-based likelihoods of the BCR sequences, AntibodyForests can be employed for integration and downstream analysis. B) An example tree colored on the heavy chain pseudolikelihood using the PLM Ablang (10), which was integrated into the AntibodyForests object using the Af_add_node_feature function. C) The correlation between the pseudolikelihood and the Levenshtein distance to the germline using the Af_distance_scatterplot function. D) The per-residue likelihood rank of the substitution along the edges of the trees using the Af_PLM_dataframe and Af_plot_PLM functions E) The average per-residue likelihood rank of the mutating and conserved residues along the edges of the trees using the Af_plot_PLM_mut_vs_cons function. F) The Jensen-Shannon divergence of the substitution rank distributions between samples using the Af_compare_PLM function. Single-cell BCR data for this analysis was derived from Kim et al. (6)

**Figure S7.**
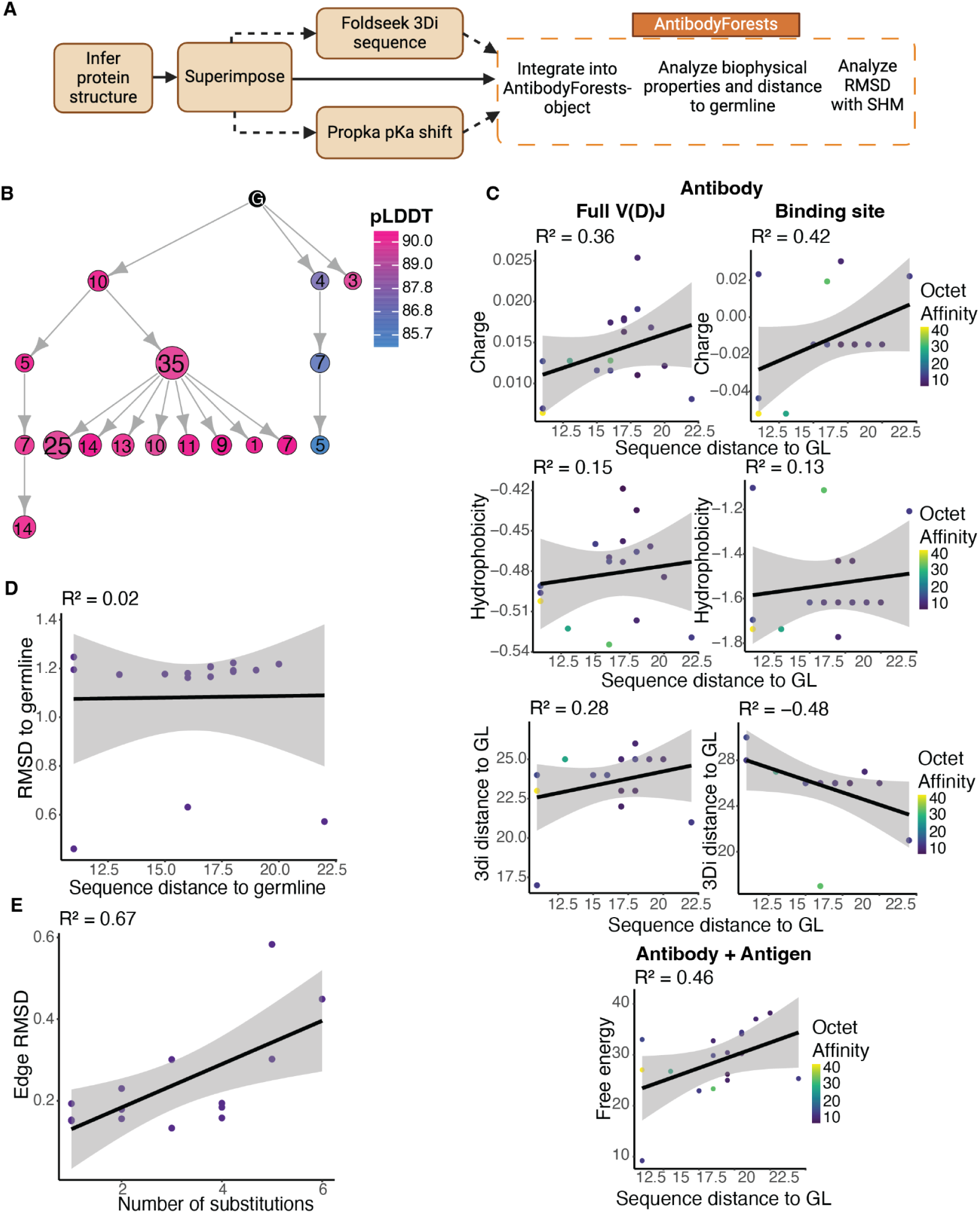
Example of how AntibodyForests can integrate protein structure evolution with sequence evolution. A) Overview of the antibody structure workflow within AntibodyForests. After inferring the 3D structure of the antibody-antigen complexes AntibodyForests can be employed for integration and downstream analysis. B) An example tree colored on the average pLDDT score provided by AlphaFold 3 (11). Lower confidence scores from AlphaFold3 were observed for a specific branch in the lineage tree, suggesting more intrinsically disordered regions. C) The correlation between biophysical properties and the Levenshtein distance to the germline for the full length V(D)J sequence (left) and for the antibody-antigen binding residues (right), and the full antibody-antigen complex (bottom). These properties were calculated with the VDJ_3d_properties function and integrated into the AntibodyForests object with the function Af_add_node_feature. D) The correlation between the root-mean-square deviation (RMSD) from the germline and the Levenshtein distance to the germline for the full length antibody generated with the VDJ_3d_properties function. E) The correlation between the RSMD and the number of substitutions over each edge in the tree generated with the Af_edge_RMSD function. Single-cell BCR and binding affinity data for this analysis were derived from Neumeier et al. (9).

## Online methods

### Data analysis

For **Figures S2**, **S5,** and **S6**, the raw BCR reads from all samples generated by Kim et al. (6) were processed with 10X Genomics’ Cell Ranger and the VDJ_build function of the Platypus package (version 3.6.0) in R (v4.4.0) (7). We applied the following parameters to the VDJ_build() function: trim.germlines = T, remove.divergent.cells = T, complete.cells.only = T. These parameters trim the germline to the V(D)J region, remove cells with more than 1 V(D)J transcript, and remove cells missing either a heavy or light chain transcript. Next, all samples per donor (retrieved at different time points after vaccination and from both the blood and lymph nodes), were reclonotyped using the Platypus function VDJ_clonotype_v3_w_enclone. We applied the following parameters to the VDJ_clonotype_v3_w_enclone() function: clone.strategy = “enclone” and global.clonotype = T. Under these parameters, the function uses 10X Genomics’ Enclone tool to group cells from all samples by V- and J-gene and CDR3 length, and shared SHM with a CDR3 nucleotide identity of at least 85%. For **Figure S5A** we only reclonotyped the time points and kept the tissues as separate clonotypes. This reclonotyped VDJ-dataframe obtained with Platypus is then supplied to AntibodyForests.

For **Figure S2**, we subset the VDJ-dataframe to only include donor 368-10. We run the function Af_build to construct lineage trees for each clonotype with the following parameters: sequence.columns = c("VDJ_sequence_aa_trimmed","VJ_sequence_aa_trimmed"), germline.columns = c("VDJ_germline_aa_trimmed","VJ_germline_aa_trimmed"), and construction.method = "phylo.tree.mp". With these parameters, nodes in the trees represent both the heavy and light chain sequence, and the maximum parsimony method is used to construct the trees, by default the Levensthein distance is used as the distance metric. To create the figures, we use the function Af_plot_tree with the following parameters: sample = "368-10", clonotype = "clonotype40", label.by = "name", edge.label = "original", show.inner.nodes = F (for subfigure A) and show.inner.nodes = T (for subfigure B). With these parameters, the lineage tree of clonotype 40 is plotted, the nodes are labeled by their assigned node names from the AntibodyForests-object, the edges are labeled by Levensthein distance between the respective nodes, and the nodes are by default colored on isotype and sized by expansion.

For **Figure S5**, we run the function Af_build on all samples with the following parameters: sequence.columns = c("VDJ_sequence_aa_trimmed","VJ_sequence_aa_trimmed"), germline.columns = c("VDJ_germline_aa_trimmed","VJ_germline_aa_trimmed"), and construction.method = "phylo.network.default". For the across-repertoire comparison we run the function Af_compare_across_repertoire with the parameters: metrics = c("spectral.density", "mean.edge.length", "betweenness"), plot = “boxplot”, significance = T. For clustering of the trees, we supply the AntibodyForests-object to the function Af_compare_within_repertoires with the following parameters: min.nodes = 20, distance.method = "jensen-shannon", clustering.method = "mediods", visualization.methods = "heatmap". With these parameters, we only consider trees with at least 20 nodes for topology analysis. We perform a k-mediods clustering on Jensen-Shannon divergence and create a heatmap. To inspect the difference in topology metrics between the clusters, we use the function Af_cluster_metrics with the parameters: clusters = output$clustering (output from Af_compare_within_repertoires), metrics = "spectral.density", min.nodes = 20, and significance = T. Next, additional node features from the VDJ-dataframe are added to the AntibodyForests-object with the function Af_add_node_feature. The influence of these parameters on the clustering is then analyzed with the function Af_cluster_node_features with the parameters: features = c("vgene_family", "timepoint", "isotype"), fill = "max", clusters = output$clustering (output from Af_compare_within_repertoires), and significance = T. These parameters take the most abundant node feature per tree in each cluster and calculates if there is a significant difference in cluster composition. Lastly, to analyze the potential difference in average distance to the germline of certain node features, we use the function Af_distance_boxplot with the parameters: distance = "edge.length", min.nodes = 10, node.feature = "timepoint" or “tissue”, groups = c("d28", "d60", "d110", "d201") or NA (when node.feature is “tissue”), significance = T, and unconnected = T. These create boxplot for the average sum of edge lengths per group per tree of at least 10 nodes and calculate the T-test p-value.

For **Figure S6**, we run the function Af_build() on all samples with the following parameters: sequence.columns = "VDJ_sequence_aa_trimmed", germline.columns = "VDJ_germline_aa_trimmed", and construction.method = "phylo.network.default". Next, we run the PLM Ablang (10) to construct probability matrices and calculate pseudolikelihoods using the code in this repository: https://github.com/dvginneken/PLM-pipeline. We use the function Af_add_node_feature to add the pseudolikelihoods to the AntibodyForests-object. The correlation between pseudolikelihood and distance to the germline was analyzed with the function Af_distance_scatterplot with the parameters: min.nodes = 5, color.by = “sample”, color.by.numberic = F, correlation = “pearson”. This creates a scatterplot of the pseudolikelihood of sequences in the trees with a least 5 nodes, the dots are categorically colored on the donor ID and a pearson correlation coefficient is calculated. Mutations along the edges of the tree were analyzed by supplying the PLM probability matrices to the function Af_PLM_dataframe and plotted with Af_plot_PLM with the parameters: group_by = “sample_id” and values = “substitution_rank”. The average ranks of the mutating and conserved residues were analyzed with the function Af_plot_PLM_mut_vs_cons with the parameters: values = "rank", dots = "all_edges", group_by = "sample_id". Finally, the PLM likelihood patterns between samples were compared with the function Af_compare_PLM.

For Figure **S3**, the preprocessed BCR reads one of the mice from Neumeier et al. (9) were processed with the VDJ_build function of the Platypus package (version 3.6.0) in R (v4.4.0) (7). We applied the following parameters to the VDJ_build() function: trim.germlines = T, remove.divergent.cells = T, complete.cells.only = T. A data frame with the bulk transcript was integrated into this VDJ-dataframe with the AntibodyForests function VDJ_integrate_bulk using the parameters: organism = “mouse”, trim.FR1 = T, and tie.resolvement = “all”. This function annotates the single-cell and bulk transcripts using IgBLAST (12), trims the FR1 regions from all the sequences and reconstructed germline sequences to account for variation in primer design, and merges the bulk transcripts into the existing single-cell clonotypes based on identical CDR3 length, V and J gene usage, and CDR3 sequence similarity. We run Af_build on the integrated VDJ-dataframe with the parameters: sequence.columns = "VDJ_sequence_aa_trimmed", germline.columns = "VDJ_germline_aa_trimmed", construction.method = "phylo.network.default", parallel = F, node.features = c("dataset"). Next, we plot clonotype 3 and 5 using the function Af_plot_tree.

For Figure **S7**, we used the data from the largest binding IgG clone of Mouse1 from Neumeier et al. (9). A lineage tree was created in R (v4.4.0) with AntibodyForests Af_build using the parameters: sequence.column = c("VDJ_sequence_aa", "VJ_sequence_aa"), germline.columns = c("VDJ_germline_aa", "VJ_germline_aa"), node.features = c("octet.affinity"). Next, we predicted the antibody-antigen complex 3D structure of the heavy and light chain of each node together with the antigen Ovalbumin (SERPINB14) using AlphaFold3 (11). The resulting structures were superimposed on the C-alpha carbon positions. 3Di sequences were determined using mini3di (v0.2.1) (13) and pKa values and free energy were computed using propka (v3.5.1) (14), both in Python (v3.11.10). Structural properties were then calculated incorporated into AntibodyForests with the function VDJ_3d_properties using the parameters: properties = c("charge", "3di_germline", "hydrophobicity", "RMSD_germline", "pKa_shift", "free_energy", "pLDDT"), chain = "HC+LC" (for S7C left and S7E,F) or chain = "whole.complex” (for S7B,C right), sequence.region = "full.sequence" (for S7C left, S7D,E,F) or “binding.residues” (for S7C right.). These structural properties were added to the AntibodyForests object using the function Af_add_node_feature and distance to the germline was plotted with Af_distance_scatterplot with parameters: correlation = "pearson", color.by = "octet.affinity", color.by.numeric = T, geom_smooth.method = "lm". Figure S7F was created with the function Af_edge_RMSD.

## Data visualization

Figures 1, **S1, S4, S6A** and **S7A** were created with Biorender.com. Lineage trees in Figures **S2**, **S3**, **S6B**, and **S7B** were generated with the function Af_plot_tree. The heatmap of the across tree construction method in Figure **S4** was generated with Af_compare_methods. The heatmap in Figure **S5B** was generated with Af_compare_within_repertoires. The boxplots in Figure **S5C** were generated with Af_cluster_metrics. The barplots in Figure **S6C** were generated with Af_cluster_node_features. The boxplots in Figure **S5E** were generated with Af_distance_boxplot. The scatterplot in Figure **S6C**, **S7C,D,E** were generated with Af_distance_scatterplot. The distribution plots in Figures **S6D** was generated with Af_plot_PLM. The boxplot in Figure **S6E** was generated with Af_plot_PLM_mut_vs_cons. The heatmap in Figure **S6F** was generated with Af_compare_PLM. The scatterplot in Figure **S7F** was generated with Af_edge_RMSD.

## Data availability

Data for Figures **S2**, **S5** and **S6** was downloaded from the NCBI Sequence Read Archive using SRA Toolkit with ID PRJNA777934 (6). Data for Figures **S3** and **S7** was provided by Neumeier et al. (9).

